# Presence of midline cilia supersedes the expression of *Lefty1* in forming the midline barrier during the establishment of left-right asymmetry

**DOI:** 10.1101/324533

**Authors:** Natalia A Shylo, Dylan Ramrattan, Scott D Weatherbee

## Abstract

Cilia in the vertebrate left-right organizer are required for the original break in left-right (L-R) symmetry. Subsequently, proper L-R patterning relies on asymmetric expression of *Nodal* in the lateral plate mesoderm (LPM). *Lefty1*, expressed in the embryonic midline, has been defined as the midline barrier, restricting the expression of *Nodal* to the left LPM. Here we use the mouse ciliary transition zone mutant *Mks1_krc_* that has left isomerism and bilateral expression of the NODAL target *Pitx2*, to reveal that the expression of *Lefty1* in the midline is insufficient for the establishment of the midline barrier. We further show through a comparison of two *Tmem107* mutants that cilia in the midline are required to supplement *Lefty1* expression and establish the functional midline barrier. *Tmem107^null^* mutants have no cilia in the midline and display left isomerism due to the loss of the midline barrier, whereas *Tmem107^schlei^* hypomorphic mutants have numerous cilia in the node and the midline, leading to normal *Lefty1* expression and L-R patterning. This study reveals a novel role for cilia in the maintenance of L-R asymmetry.

## INTRODUCTION

Heterotaxy, an abnormal arrangement of internal organs along the left-right (L-R) axis, typically involves severe cardiac abnormalities, associated with a high mortality rate (Kim, 2011; Lin, *et al*, 2014). Mouse models have been essential to discern the function of genes mutated in patients and identify new heterotaxy loci (Li, *et al*, 2015; Norris and Grimes, 2012). Among known heterotaxy-associated genes many are involved in cilia formation and function (Li, Klena, Gabriel, *et al*, 2015; Pennekamp, *et al*, 2015). Motile cilia in the left-right organizer (node in mouse, gastrocoel roof plate in Xenopus, Kupffer’s vesicle in zebrafish) create a leftward flow of extraembryonic fluid that breaks bilateral L-R symmetry. Mutations that lead to paralyzed cilia, as in *Dnah11^iv^* mutant mice, or as seen in patients with primary ciliary dyskinesia, result in randomized L-R patterning, including left and right isomerism – left or right bilaterally symmetric organs (Dougherty, *et al*, 2016; Layton, 1976; Piedra, *et al*, 1998; Seo, *et al*, 1992; Supp, *et al*, 1997). Sensation of flow by cilia on the crown cells of the node is necessary for degradation of *Cerl2* mRNA in the crown cells at the left of the node, which relieves *Cerl2* inhibition of *Nodal* expression in these cells and ultimately leads to the induction of the left-side determinant *Nodal* in the left lateral plate mesoderm (LPM) (Marques, *et al*, 2004; Yoshiba, *et al*, 2012). Mice harboring mutations in the ciliary-localized calcium channel Polycystin2 (PC2, *Pkd2* gene) fail to induce *Nodal* in the left LPM due to their inability to sense flow (Yoshiba, Shiratori, Kuo, *et al*, 2012). Without this “leftness” signal, animals develop two right sides (Pennekamp, *et al*, 2002).

NODAL is a secreted TGF-TGF-β family member, capable of autoactivation. The NODAL cascade is a signal transduction pathway that propagates L-R asymmetric patterning to other tissues and organs. Similar to mutants with a loss of flow sensation, mouse mutants that fail to transduce *Nodal* signal in the left LPM also develop right isomerism (Saijoh, *et al*, 2003; Yan, *et al*, 1999). Diffusible NODAL is believed to be restricted to the left side of the embryo by a midline barrier, mediated by LEFTY1 - also a secreted TGF-β family member - which can bind to NODAL, thus preventing its diffusion across the midline and its activation of the left-sided Nodal cascade on the right side of the embryo (Chen and Shen, 2004; Meno, *et al*, 1998). Mouse *Lefty1* mutants have left isomerism, thought to be secondary to an abnormal midline barrier (Meno, Shimono, Saijoh, *et al*, 1998). Together, this system ensures that the initial symmetry break at the node is propagated to and maintained within the rest of the embryo.

Mounting evidence suggests that cilia have roles in L-R patterning beyond inducing and sensing the flow at the node. For example, *Kif3a*, *Ift172*, *Ift88* and *Ift38* mutants don’t form cilia but display bilateral marker expression in the LPM, while current models would suggest that loss of cilia should result in a failure to generate and sense the flow, and induce the NODAL cascade, resulting in right isomerism (Botilde, *et al*, 2013; Grimes, *et al*, 2016; Huangfu, *et al*, 2003; Murcia, *et al*, 2000; Takeda, *et al*, 1999). Furthermore, loss of cilia is epistatic to the loss of *Pkd2*, as evidenced by left isomerism in *Kif3a^null^*; *Pkd2^null^* embryos (Grimes, Keynton, Buenavista, *et al*, 2016), indicating a role for cilia downstream of flow sensation by Pkd2.

Previously described mouse *Tmem107^schlei^* and human patients with mutations in *Tmem107*, which encodes a ciliary transition zone protein, display fewer and abnormally shaped cilia, but no L-R patterning defects (Lambacher, *et al*, 2016; Shaheen, *et al*, 2015; Shylo, *et al*, 2016; Weatherbee, *et al*, 2009). In this study we report that *Tmem107^null^* mice have left isomerism, resembling cilia mutants with global loss of cilia. We took advantage of the *Tmem107* mutants to examine additional roles for cilia in establishing the L-R axis. We show that while *Tmem107^null^* mutants have a randomized *Nodal* expression at the node, they develop bilateral *Nodal* expression in the LPM, which underlies left pulmonary isomerism. Using an allelic series of *Tmem107* mutations, and an additional transition zone mutant, *Mks1_krc_*, that also has few cilia and left pulmonary isomerism (Weatherbee, Niswander and Anderson, 2009), we uncover an unexpected but essential role for cilia in establishment and maintenance of the midline barrier, independent of *Lefty1* expression.

## RESULTS

### *Tmem107^null^* mutants display multiple left-right defects

*Tmem107* encodes a ciliary transition zone protein, required for the formation of a functional transition zone in primary cilia (Lambacher, Bruel, van Dam, *et al*, 2016; Shylo, Christopher, Iglesias, *et al*, 2016). Given the ubiquitous nature of primary cilia, we were surprised to see an enrichment of *Tmem107* mRNA in the embryonic node at E7.5-8.5, compared to all other embryonic tissues (Fig. 1 A, B and Fig. S1), suggesting a possible role for TMEM107 in the node at this early stage. This expression pattern also appeared at odds with the lack of L-R abnormalities in a previously described ENU-induced *Tmem107^schlei^* mouse mutant (Christopher, *et al*, 2012). However, close examination of the *Tmem107^tm1Lex^* mutant, which completely lacks the *Tmem107* gene product (Tang, *et al*, 2010), revealed multiple heterotaxy characteristics, including randomized heart looping and stomach positioning (Fig. 1 C-F). These data indicate that TMEM107 is critical for establishing the L-R axis and that *Tmem107^schlei^* is a hypomorphic allele. To determine the extent to which TMEM107 regulates L-R patterning we further examined the heterotaxy phenotype in *Tmem107^tm1Lex^* (hereafter referred to as*Tmem107^null^*) mutants.

**Figure 1.**
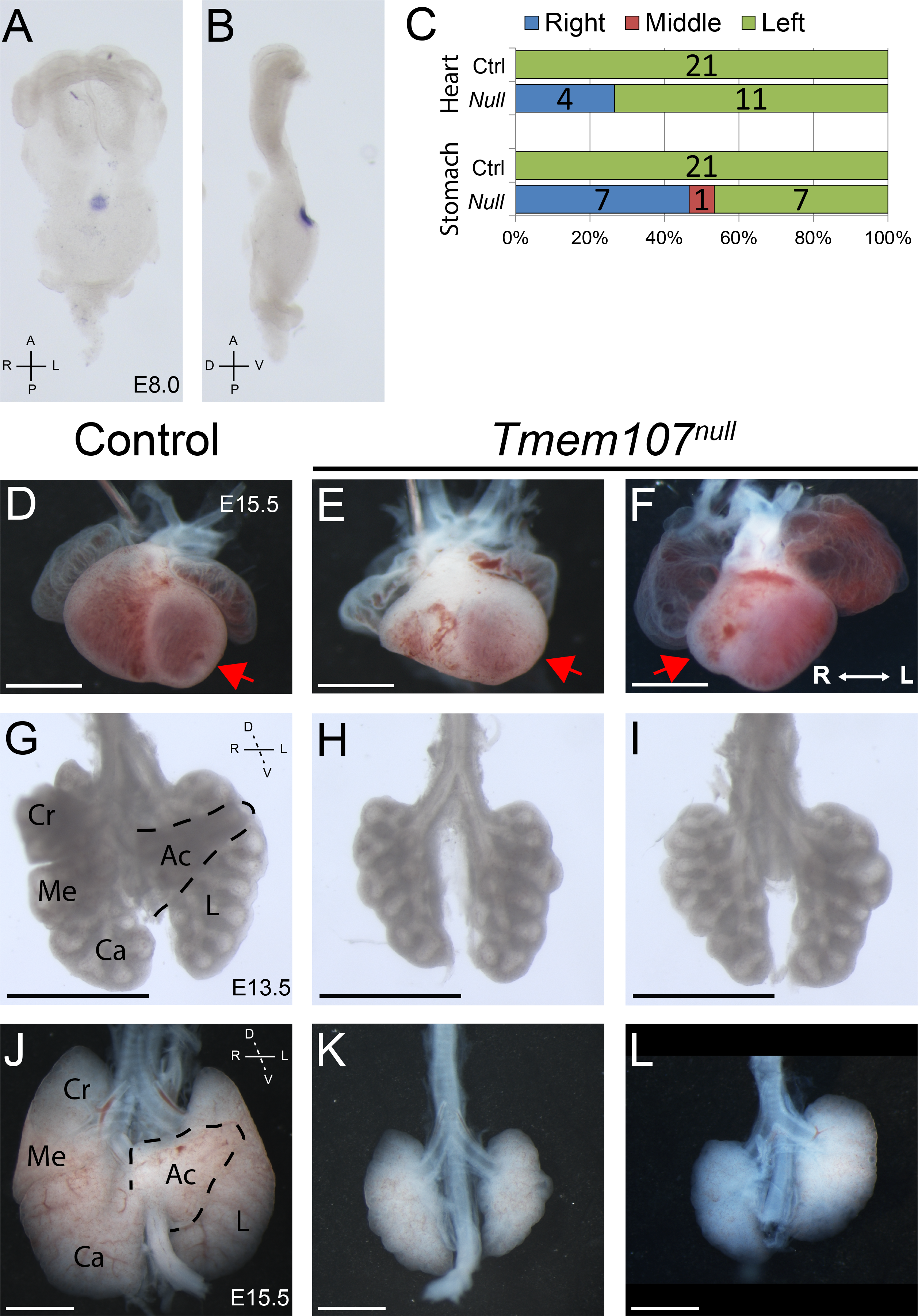
*Tmem107^null^* mutants display multiple left-right patterning defects. (A, B) *Tmem107* mRNA is strongly enriched in the nodes of E8.0 embryos, as shown in (A) ventral view and (B) lateral view. (C) Heart positioning (based on the orientation of the apex to the left or right) and stomach positioning are randomized in *Tmem107^null^* embryos. Numbers represent individual animals. (D-F) Ventral view of E15.5 hearts. Control hearts have their apices on left (D, red arrow), while *Tmem107^null^* hearts have randomized looping with the apex on left (E, arrow) or right (F, arrow). (G-L) Ventral view of E13.5 (G-I) and E15.5 (J-L) lungs. Control mouse lungs have 4 right lobes and a single left lung lobe, while *Tmem107^null^* lungs display single lobes on both sides (H, I, K, L), suggesting left pulmonary isomerism. Pulmonary hypoplasia in *Tmem107^null^* mutants becomes pronounced by E15.5 (K, L). Cr - cranial lobe, Me - medial lobe, Ca - caudal lobe, Ac - accessory lobe, L - left lobe. Scale bars = 1 mm.

The position of the heart apex was used to determine directionality of heart looping and reveal its randomization. 4 out of 15 *Tmem107^null^* mutants showed clear, abnormal right-side looping of the heart (Fig. 1C-F, red arrows in D-F). Stomach positioning appeared to be independent of heart directionality and showed perfect randomization in *Tmem107^null^* embryos compared to the normal left-sidedness (Fig 1C). We also examined the developing lungs, which, in control mice, form one lobe on the left side, and four lobes on the right side (Fig. 1G, J). The lungs in *Tmem107^null^* embryos were bilaterally single-lobed (Fig. 1H, I, K, L). Careful examination of E13.5 lungs revealed that both single lobes had equivalent numbers of branching points, consistent with normal branching pattern for the left lobe at that stage (Metzger, *et al*, 2008), leading us to conclude that *Tmem107^null^* embryos develop left pulmonary isomerism (Fig. 1 G-I). At later stages, *Tmem107^null^* lungs become severely hypoplastic, however in all cases examined, only a single lobe was observed on both sides (Fig. 1K-M).

### L-R perinodal signaling is disrupted in *Tmem107^null^* mutants

Left-right patterning of the organs begins at the node, and the earliest marker of the L-R symmetry break is degradation of *Cerl2* mRNA on the left side of the node in response to cilia-generated flow (Inacio, *et al*, 2013; Kawasumi, *et al*, 2011; Marques, Borges, Silva, *et al*, 2004; Nakamura, *et al*, 2012; Schweickert, *et al*, 2010; Shinohara, *et al*, 2012). Left-side *Cerl2* degradation is apparent by 2-3 somites stage in controls and becomes more pronounced at the 4-5 somites stage (Fig. 2A, B, E). In contrast, *Cerl2* expression in *Tmem107^null^* embryos is strong bilaterally at the 2-3 somites stage, and either maintains strong bilateral expression, or becomes degraded in a random fashion by 4-5 somites stage (Fig. 2 C, D, F). Normally, left-sided *Cerl2* degradation alleviates its inhibition of *Nodal*, and leads to subsequent *Nodal* upregulation on the left side of the node (Fig. 2 G-K). We expected that the strong bilateral *Cerl2* expression would lead to bilateral down-regulation of *Nodal* (Marques, Borges, Silva, *et al*, 2004), but we observed randomized *Nodal* expression levels around *Tmem107^null^* nodes (Fig. 2I-L). These results suggest a disconnect between the expression of Cerl2 and Nodal in *Tmem107^null^* embryos.

**Figure 2.**
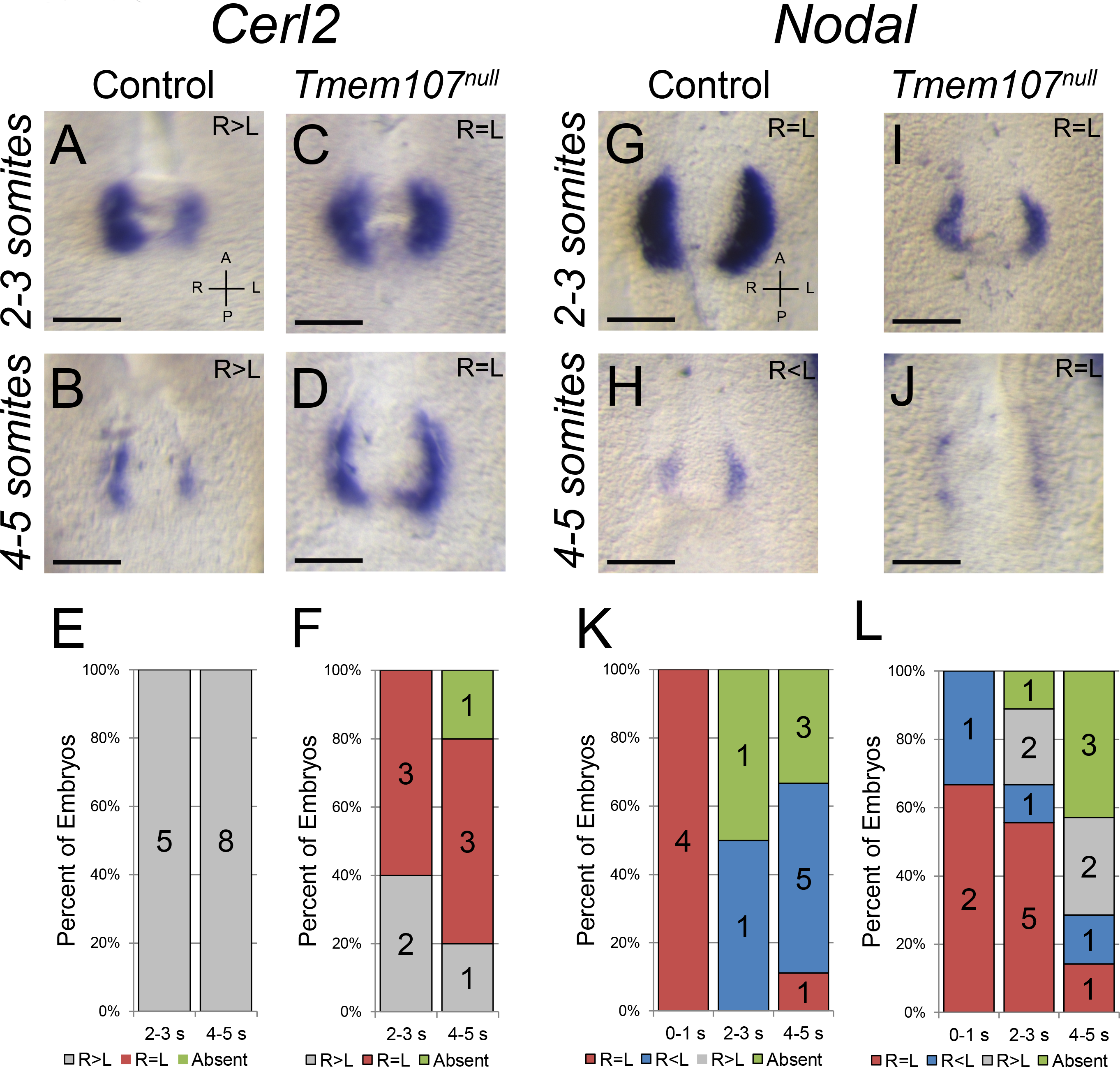
Flow-induced signaling in *Tmem107^null^* nodes is compromised. (A-D) *in situ* hybridization for *Cerl2*, at early (2-3 somites) and late (4-5 somites) stages. Control embryos (A, B) show progressive degradation of *Cerl2* mRNA on the left side while *Tmem107^null^* embryos (C, D) maintain strong bilateral *Cerl2* mRNA, with some randomization of expression at later stages. (G-J) *in situ* hybridization for *Nodal*, at early (2-3 somites) and late (4-5 somites) stages. Control embryos (G, H) show reduced *Nodal* expression on the right side at late stages (H), while *Tmem107^null^* embryos (I, J) maintain bilateral *Nodal* mRNA for longer, and ultimately have a randomized pattern of expression (L). (E, F, K, L) Graphs accompanying each probe represent the quantification of the embryos with a given expression pattern, broken down by somite number. Scale bars are 100 μm.

### Cilia in the nodes of *Tmem107^null^* embryos maintain their identities

*Cerl2* degradation is dependent on the ability to sense nodal flow (Nakamura, Saito, Kawasumi, *et al*, 2012; Schweickert, Vick, Getwan, *et al*, 2010), so the lack of early left-sided *Cerl2* degradation points to abnormal motile cilia in the pit of the node or flow sensing cilia surrounding the node. We used ARL13B as a marker of cilia, and the nodes of E8.0 *Tmem107^null^* embryos were the only region where we detected ARL13B+ cilia, as compared to extensive ARL13B+ cilia throughout the control embryo (Fig. 3 A-D). Generally, we observed fewer ARL13B+ cilia in *Tmem107^null^* mutant nodes when compared to controls (Fig. 3A-D), which is consistent with cilia number reductions reported in *Tmem107^schlei^* mouse mutants, and in human patients with mutations in *TMEM107*. (Christopher, Wang, Kong, *et al*, 2012; Lambacher, Bruel, van Dam, *et al*, 2016; Shaheen, *et al*, 2013; Shylo, Christopher, Iglesias, *et al*, 2016). The loss of ARL13B+ cilia in the endoderm, surrounding the node (Fig. 3 A-F) stands in contract with the specific enrichment of *Tmem107* mRNA in the node only. These data indicate that TMEM107 is required throughout the embryo and, in addition to its specific enrichment at the node (Fig. 1 A, B, S1), *Tmem107* must be expressed in other tissues, but below the detection levels of *in situ* hybridization.

**Figure 3.**
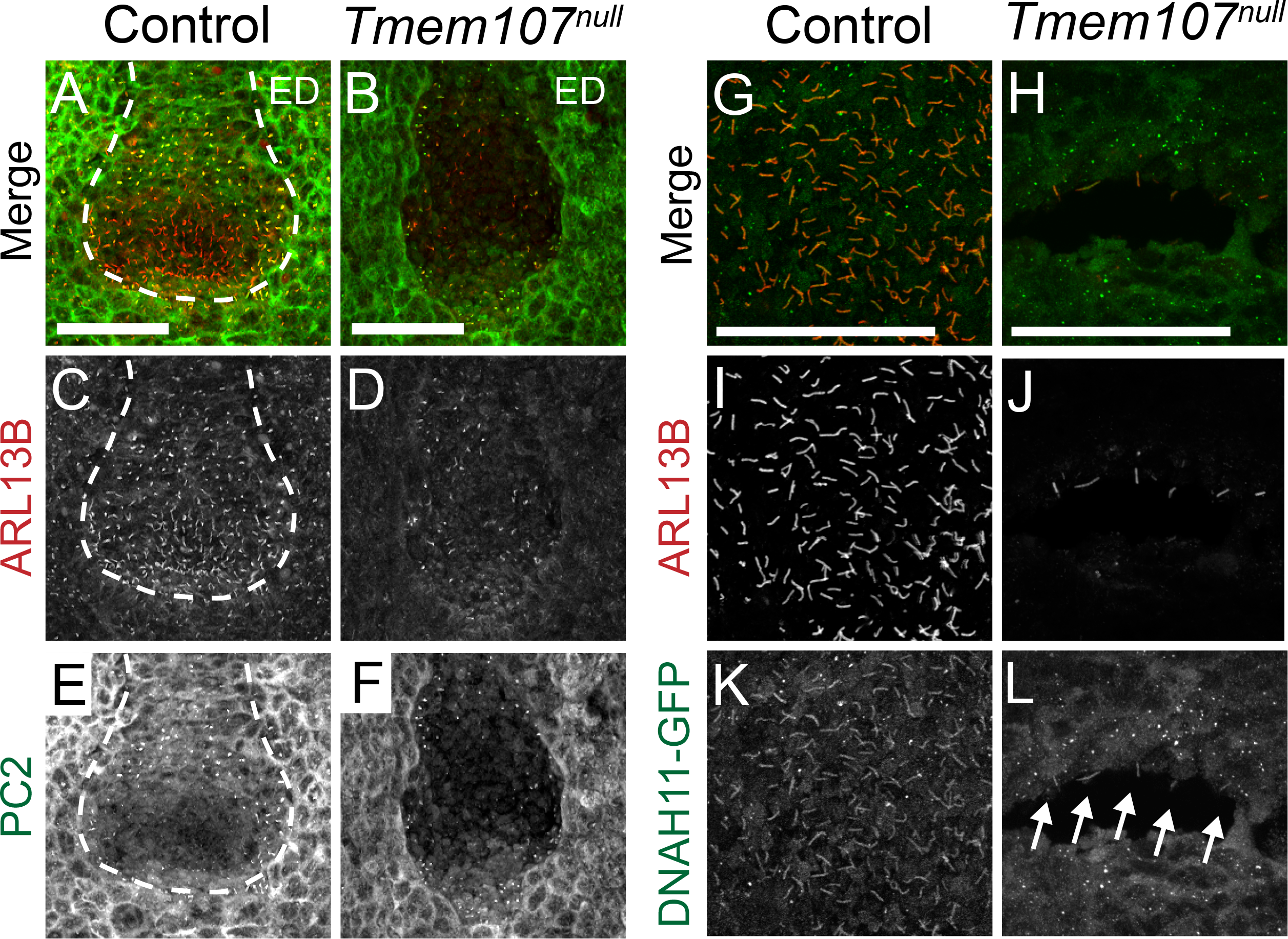
Motile and sensory cilia are present in *Tmem107^null^* nodes, but in reduced numbers. (A-L) Maximum intensity projected Z-stacks of confocal immunofluorescent images of E8.0 nodes. (A, C) Low magnification images of control nodes show ARL13B (red) present in all cilia in the node and the surrounding cells. (B, D) In *Tmem107^null^* embryos ARL13B is only present in a small population of nodal cilia, and not in the surrounding cells. (A, E) Polycystin2 (PC2, green) is enriched in sensory cilia on crown cells, around the edge of the node. The distribution of PC2 positive cilia appears normal in *Tmem107^null^* nodes (B, F). (G-L) Higher magnification views of the nodal pits of control and *Tmem107^null^* embryos that are also homozygous for the *Dnah11^GFP^* transgene. ARL13B (I, J, and red in G, H) marks the nodal cilia showing that the nodal pit of *Tmem107^null^* embryos has only a small population of ARL13B positive cilia. Control embryos have DNAH11-GFP positive motile cilia (G, green and K) throughout the pit of the node. Cilia in the pit of *Tmem107^null^* nodes also have motile identity as marked by DNAH11-GFP (H, green and L). ED - endoderm. Scale bars are 50 μm.

Very few motile cilia in the node pit are necessary to generate enough flow to establish proper L-R patterning (Shinohara, Kawasumi, Takamatsu, *et al*, 2012). To test whether *Tmem107^null^* mutants lack cilia with motile identity specifically, we introgressed a transgenic reporter for an axonemal dynein required for ciliary motility, *Dnah11*, fused to *GFP* (*Dnah11-GFP*) (McGrath, *et al*, 2003). All cilia in the pit of control nodes are DNAH11-GFP+ (Fig. 3 G, I, K). We found that most cilia in the pit of *Tmem107^null^* nodes are also DNAH11-GFP+ (Fig. 3 H, J, L) indicating that motile cilia are specified in the absence of TMEM107 function. To examine whether the *Tmem107^null^* nodes have “sensing” cilia, we next examined the localization of PC2, which is enriched in crown cells surrounding the node and is required for flow sensing (Yoshiba, Shiratori, Kuo, *et al*, 2012) We observed many PC2+ cilia in *Tmem107^null^* crown cells, suggesting that ciliated cells with flow-sensing identity are preserved in the mutants (Fig 3 E, F). Thus, nodes of *Tmem107^null^* embryos maintain both cilia with motile and sensory identities, albeit at largely decreased numbers.

### *Nodal* and *Pitx2* are expressed bilaterally in the lateral plate mesoderm of *Tmem107^null^* embryos

The randomized *Nodal* expression in *Tmem107^null^* nodes (Fig. 2 G-L) was inconsistent with the left pulmonary isomerism that we observed in older embryos (Fig. 1 H-M). Asymmetric *Nodal* at the node is thought to signal to the lateral plate mesoderm (LPM) and autoactivate robust *Nodal* expression specifically in the left LPM (Fig 4. A, B) (Kawasumi, Nakamura, Iwai, *et al*, 2011; Yamamoto, *et al*, 2003). *Nodal* expression in the left LPM is transient, starting around the 2-3 somites stage, and becomes extinguished a few hours later (6-7 somites stage). To test if there were defects in the transmission of the NODAL signal from the node to the LPM, we examined *Nodal* LPM expression between 0-5 somites (Fig. 4 A, B). Consistent with the lung phenotype, the vast majority of *Tmem107^null^* embryos displayed bilateral *Nodal* transcripts in the LPM (Fig. 4 A, B) by 4-5 somites stage. These data are consistent with previous reports of dissociation between *Nodal* expression at the node and its expression in the LPM in other mutants, although the mechanism by which this occurs is unclear. (Ishimura, *et al*, 2008; Marques, Borges, Silva, *et al*, 2004; Norris, *et al*, 2002).

**Figure 4.**
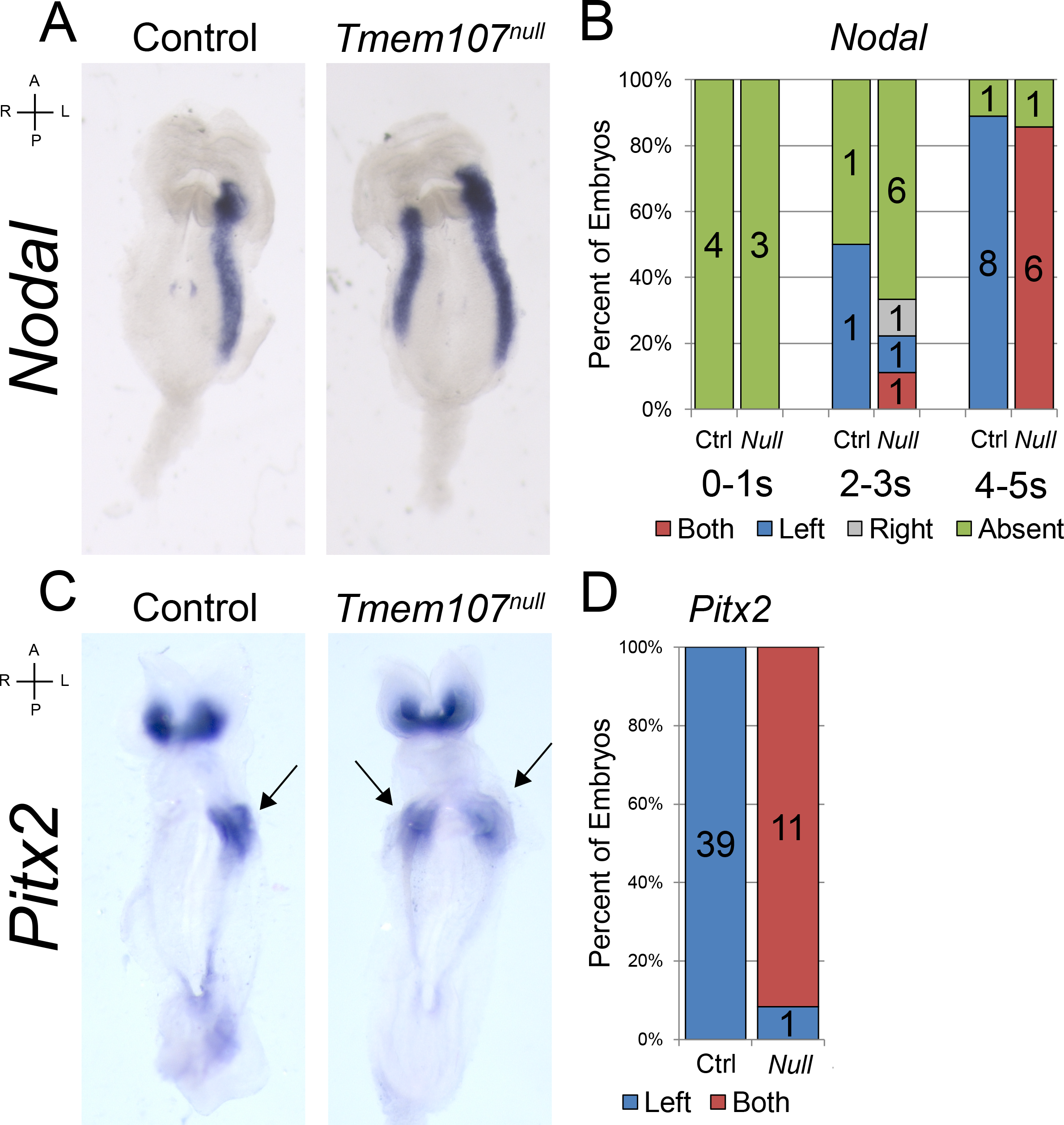
The lateral plate mesoderm in *Tmem107^null^* embryos acquires bilateral left identity. (A) *in situ* hybridization for *Nodal* in representative control and *Tmem107^null^* E8.0 embryos showing left side restriction in controls and bilateral expression in mutants. (B) Quantification of the expression patterns for *Nodal* at the LPM, broken down by somite stage. (C) *in situ* hybridization for *Pitx2* in representative control (left LPM, arrow) and *Tmem107^null^* (bilateral, arrows) E8.5 embryos. *Pitx2* is normally expressed bilaterally in the developing heads of control and mutant embryos. (D) Quantification of the *Pitx2* expression localization. Numbers in all graphs indicate individual animals.

In the left LPM, NODAL signaling induces the expression of *Pitx2*, a transcription factor, whose left-side expression persists in various LPM-derived tissues, long after *Nodal* expression is lost (Fig. 4 C, D) (Lin, *et al*, 1999; Meno, Shimono, Saijoh, *et al*, 1998; Ryan, *et al*, 1998; Yoshioka, *et al*, 1998). We observed bilateral expression of *Pitx2* in the LPM of *Tmem107^null^* embryos, indicating that once left-side identity is established in the LPM by NODAL, it is transduced accordingly (Fig. 4 C, D). Together, the *Nodal* and *Pitx2* expression patterns are consistent with the left pulmonary isomerism that we observe in older *Tmem107^null^* embryos (Fig. 1 G-L), but contrast with the randomized node expression of *Nodal*.

Thus far, our data indicate that TMEM107 may have multiple functions in establishing L-R asymmetry. Despite the presence of cilia with both motile and sensory identities, we observe a failure to properly degrade *Cerl2* in crown cells. Furthermore, *Nodal* and *Cerl2* expression patterns in crown cells are inconsistent with respect to each other, suggesting that TMEM107 may potentially have roles in these early processes. Lastly, randomized *Nodal* at the node shows inconsistencies with bilateral *Nodal* at the LPM that ultimately leads to the left pulmonary isomerism. Since bilateral *Nodal* in the LPM is the most downstream effect of *Tmem107* loss on L-R development, we decided to focus our attention specifically on this aspect of L-R development.

### *Tmem107^null^* embryos have a defective barrier and disrupted SHH signaling in the midline

Consistently bilateral *Nodal* and *Pitx2* in the LPM of *Tmem107^null^* embryos led us to examine additional factors that regulate L-R identity. The midline barrier model postulates that *Lefty1*, a TGFβ family member expressed in the left prospective floorplate (Meno, *et al*, 1997), is required to inhibit the passage of NODAL to the right side of the embryo, preventing NODAL from autoactivating its expression in the right LPM (Meno, Shimono, Saijoh, *et al*, 1998; Yamamoto, Mine, Mochida, *et al*, 2003). In contrast to control embryos, none of the *Tmem107^null^* mutants expressed *Lefty1* in the midline (Fig. 5 B, C, arrowheads). The absence of *Lefty1* expression, in conjunction with bilateral *Nodal* expression in the LPM indicates the loss of a functional midline barrier. As *Lefty1* expression has been shown to be dependent on SHH signaling (Tsiairis and McMahon, 2009; Tsukui, *et al*, 1999), we examined SHH signaling in the midline of *Tmem107^null^* embryos.

**Figure 5.**
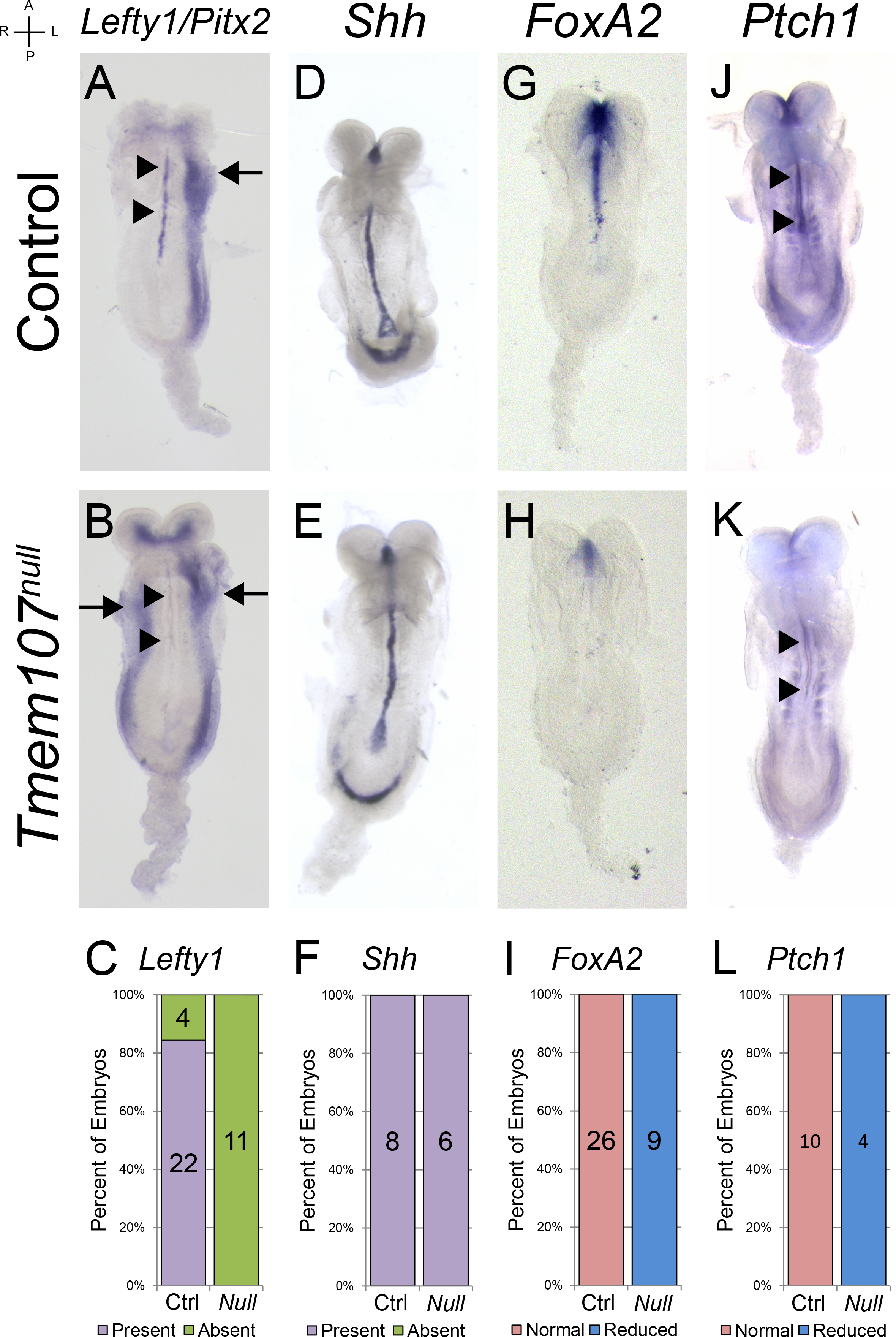
*Tmem107^null^* embryos show *Shh*-dependent loss of the midline barrier. (A, B) Double *in situ* hybridization for *Lefty1* (midline, arrowheads) and *Pitx2* (LPM, arrows) in representative control (A) and *Tmem107^null^* (B) embryos. *Pitx2* riboprobe was used as a positive control for *in situ* hybridization reaction conditions, and is expressed in left LPM in control embryos (A, arrow) but is bilateral in *Tmem107^null^* embryos (B, arrows). In contrast, *Lefty1* signal is lost in the midlines of *Tmem107^null^* embryos (B, arrowheads). (D, E) *in situ* hybridization revealed unchanged midline expression of *Shh* in *Tmem107^null^* (E) embryos compared to controls (D). (G, H) However, *in situ* hybridization for *FoxA2*, a direct target of SHH signaling, uncovered a strong reduction of midline *FoxA2* expression in *Tmem107^null^* (H) embryos compared to controls (G). (J, K) *in situ* hybridization for *Ptch1* shows midline and somite expression in control embryos (J) but a strong reduction in *Ptch1* midline signal in *Tmem107^null^* embryos (K, arrowheads). (C, F, I, L) Graphs accompanying each probe represent the quantification of the embryos with a given expression pattern.

Hedgehog signaling is tightly linked to cilia in vertebrates, and our previous studies showed reduced SHH signaling in the neural tubes of *Tmem107^schlei^* mutants (Christopher, Wang, Kong, *et al*, 2012). At E8.0, *Shh* is expressed in the node and the notochordal plate. (Fig. S2 A-D) (Echelard, *et al*, 1993; Zhang, *et al*, 2001) and SHH signaling is necessary to specify the floor plate and establish the midline (Chiang, *et al*, 1996). We did not observe any difference in midline *Shh* expression *Tmem107^null^* embryos (Fig. 5D-F, Fig. S2 A, B) and decided to look downstream of the ligand at the expression of two direct targets of SHH signaling, *FoxA2* (Fig. 5 G-I) and *Ptch1* (Fig. 5 J-L), which are both enriched in control midlines. Despite the normal expression of *Shh*, the midline enrichment of both *FoxA2* and *Ptch1* was strongly reduced in all *Tmem107^null^* embryos examined (Fig. 5 H, I, K, L, arrowheads). These results indicate that TMEM107 is required for the transduction of the midline SHH signal, to establish *Lefty1* expression and midline barrier function. These results were consistent with the current model in the field.

### Analysis of hypomorphic*Tmem107^schlei^* embryos with normal L-R patterning reveals SHH-independent expression of *Lefty1*

We previously reported that the hypomorphic *Tmem107^schlei^* mutants did not show any disruptions in L-R asymmetry of late organs, but displayed abnormal SHH signaling in limbs and neural tubes (Christopher, Wang, Kong, *et al*, 2012). Consistent with this, the most downstream marker of left-side patterning, *Pitx2*, is restricted to the left LPM of *Tmem107^schlei^* mutants, similar to control embryos (Fig. 6 A-C, J, K). *Shh* was also expressed normally in the node and midline region of all *Tmem107^schlei^* embryos examined (Fig. 6 D-F, S1C, D). In contrast, *FoxA2* was lost from *Tmem107^schlei^* mutant midlines, similar to *Tmem107^null^* embryos, and consistent with abnormal SHH signaling observed in other *Tmem107^schlei^* tissues (Fig. 6 G-I) (Christopher, Wang, Kong, *et al*, 2012). Notably, we detected consistent *Lefty1* expression in the midline region of *Tmem107^schlei^* embryos (Fig. 6 J-L). These data indicate that the mutant TMEM107 protein produced by the *Tmem107^schlei^* hypomorphic allele is sufficient for the expression of *Lefty1*, but not *FoxA2*, and is capable of establishing a midline barrier and preserving L-R identity.

**Figure 6.**
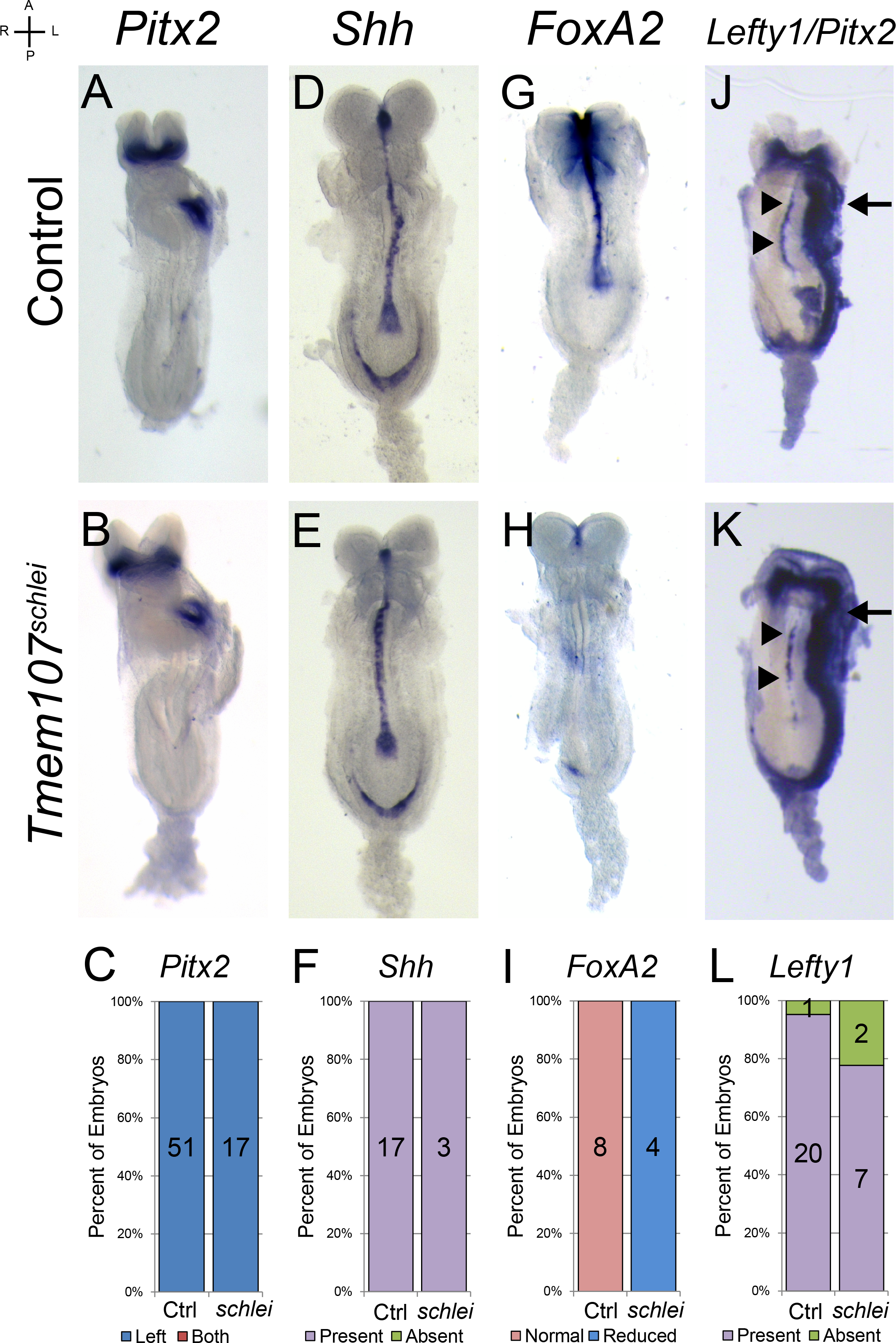
*Tmem107^schlei^* embryos maintain left-right asymmetry despite reduced *Shh* signaling in the midline. (A, B) *in situ* hybridizations for *Pitx2* in control (A) and *Tmem107^schlei^* (B) embryos show normal expression in the left LPM (D, E) *in situ* hybridization for *Shh* in control (D) and *Tmem107^schlei^* (E) embryos, showing normal midline expression in both cases. (G, H) *FoxA2 in situ* hybridization is normal in controls (G) but is strongly reduced in the midlines of *Tmem107^schlei^* (H) embryos. (J, K) Double *in situ* hybridization for *Lefty1* (midline, arrowheads) and *Pitx2* (LPM, arrows) revealed *Lefty1* expression in the midlines of *Tmem107^schlei^* (K, arrowheads) embryos, albeit faint and/or patchy. (C, F, I, L) Graphs accompanying each probe represent the quantification of the embryos with a given expression pattern.

### *Lefty1* is expressed in *Mks1^krc^* transition zone mutants with midline barrier defects

Several mouse lines carrying mutations in genes encoding transition zone proteins develop left pulmonary isomerism and/or bilateral expression of markers in the LPM (Abdelhamed, *et al*, 2015; Garcia-Gonzalo, *et al*, 2011; Vierkotten, *et al*, 2007). To test whether the midline barrier defects observed in *Tmem107^null^* embryos were specific to *Tmem107* function, or are a general feature of transition zone mutants, we examined L-R patterning in *Mks1_krc_* embryos. MKS1 is a transition zone protein that directly interacts with TMEM107, and the localization of MKS1 and TMEM107 to the transition zone of the cilium have been shown to be co-dependent in *C.elegans* (Lambacher, Bruel, van Dam, *et al*, 2016). The *Mks1_krc_* mutation is considered null, (Cui, *et al*, 2011; Weatherbee, Niswander and Anderson, 2009) and both *Mks1_krc_* and *Tmem107^null^* mutants die around E16.5 with an overlapping set of characteristics, including randomized heart looping, hypoplastic lungs, and left pulmonary isomerism (Weatherbee, Niswander and Anderson, 2009). In contrast to *Tmem107*, *Mks1* mRNA is detected in an even, ubiquitous pattern throughout the embryo, with a slight enrichment in the midline (Fig. 7 A, B). We observed bilateral LPM expression of *Pitx2* in *Mks1_krc_* mutants (Fig 7 C-E, I, J, arrows), similar to *Tmem107^null^* embryos and consistent with the observed left pulmonary isomerism. Like in *Tmem107^null^* embryos, SHH signaling in the midline of *Mks1_krc_* embryos is defective, since *FoxA2* failed to be induced in the midline (Fig. 7 F-H). Strikingly, *Lefty1* expression was detected in most of the *Mks1_krc_* embryos examined, yet bilateral *Pitx2* expression indicates that the midline barrier fails to function in these mutants (Fig. 7 I-K). These results go against the current model in the field for the establishment of the midline barrier. Similar to our results in *Tmem107^schlei^* embryos, we see *Lefty1* expression despite defective SHH signaling, suggesting that *Lefty1* expression is SHH independent. Most strikingly, though, our data show that *Lefty1* expression in *Mks1_krc_* embryos is insufficient for the establishment of the midline barrier.

**Figure 7.**
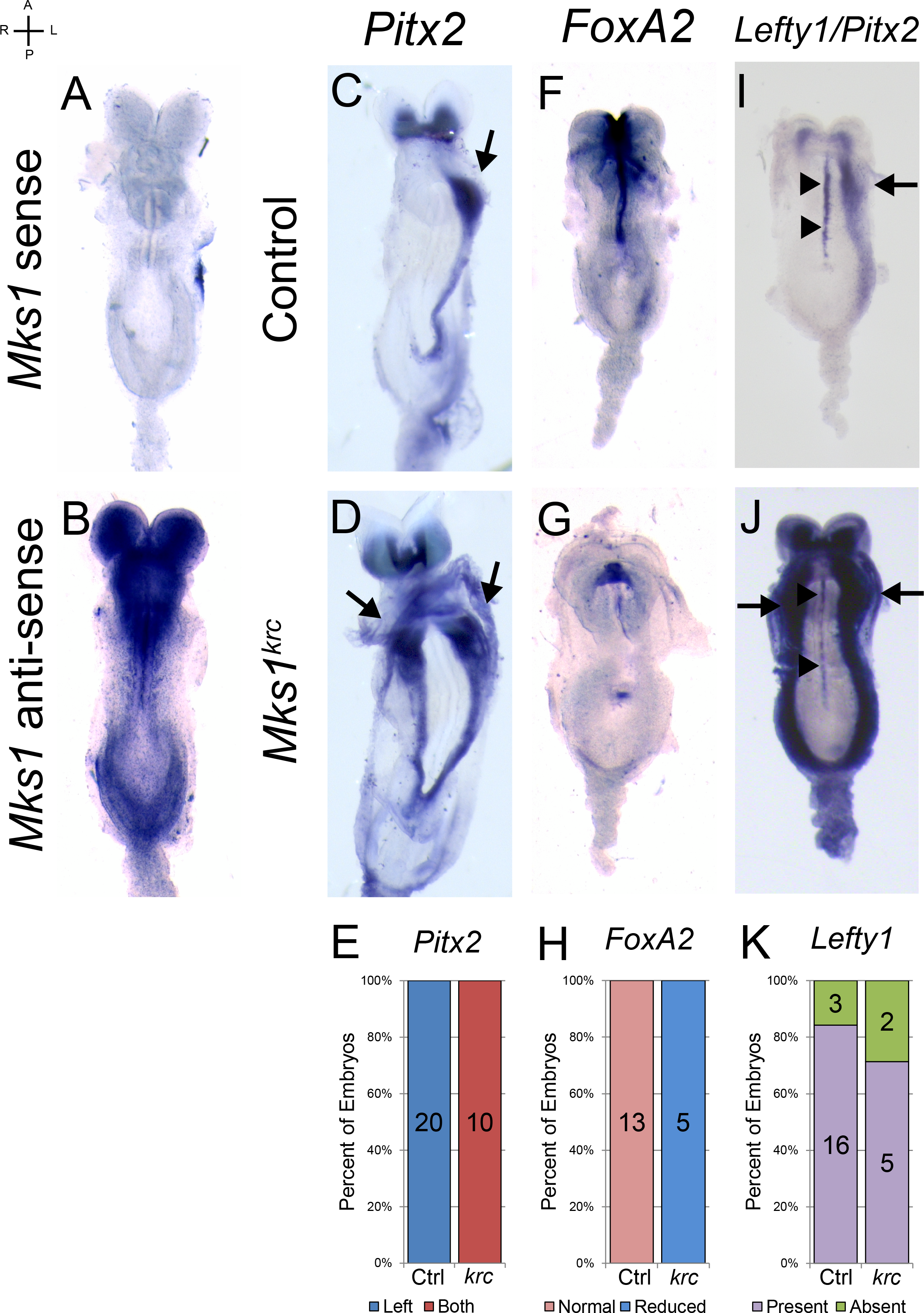
*Mks1^krc^* left-right patterning defects show similarities and differences to *Tmem107^null^*. (A, B) *Mks1 in situ* hybridizations in E8.0 control embryos revealed no clear signal for the *Mks1* sense probe (A), while the *Mks1* anti-sense probe (B) indicated that *Mks1* mRNA is present throughout the embryo, with some enrichment in the midline. (C, D) *in situ* hybridization shows that *Pitx2* expression is normal in controls (C), but bilateral in the LPM of *Mks1_krc_* embryos (D). (F, G) *FoxA2 in situ* hybridization shows strong midline expression in controls (F) but *FoxA2* expression is strongly reduced in the midlines of*Mks1_krc_* (G) embryos. (I, J) Double *in situ* hybridization for *Lefty1* (midline, arrowheads) and *Pitx2* (LPM, arrows) in embryos with 2-5 somites revealed that *Lefty1* signal is retained in the midlines of *Mks1_krc_* embryos (J, arrowheads) but is insufficient to prevent left isomerism. *Pitx2* probe was used as a positive control for *in situ* hybridization reaction conditions (I, J arrows). (E, H, K) Graphs accompanying each probe represent the quantification of the embryos with a given expression pattern.

### Midline cilia are necessary to establish midline barrier function

TMEM107 and MKS1 are both required for normal cilia number and ciliary protein localization (Christopher, Wang, Kong, *et al*, 2012; Cui, Chatterjee, Francis, *et al*, 2011; Lambacher, Bruel, van Dam, *et al*, 2016; Shaheen, Faqeih, Alshammari, *et al*, 2013; Shylo, Christopher, Iglesias, *et al*, 2016; Weatherbee, Niswander and Anderson, 2009). The abnormal midline gene expression patterns in *Mks1_krc_* and *Tmem107^null^* mutants suggest MKS1 and TMEM107 may be required for cilia formation or function in the midline. Prior studies by scanning electron microscopy (SEM) have shown loss of cilia in the nodes of *Mks1* mutants (Cui, Chatterjee, Francis, *et al*, 2011; Weatherbee, Niswander and Anderson, 2009), but normal cilia numbers in the nodes of *Tmem107^schlei^* embryos (Figure S3 and (Christopher, Wang, Kong, *et al*, 2012)). However, *Tmem107* or *Mks1* mutant midlines have not been analyzed at E8.0 for the presence of cilia.

Examination of the ARL13B cilia marker in *Tmem107^null^* embryos revealed very few midline cilia (Figure 8 B) compared to controls (Figure 8 A), consistent with reports that node cells migrate into the midline and contribute to the notochordal plate (Lee and Anderson, 2008; Sulik, *et al*, 1994). Although we observed very few cilia in the nodes of *Tmem107^null^* embryos (Fig. 3), their almost complete absence in the midline was surprising. Unlike *Tmem107^null^* mutants, *Tmem107^schlei^* embryos appear to have a near-normal numbers of cilia in their nodes (Figure S3 and (Christopher, Wang, Kong, *et al*, 2012)) and the midlines of these animals contain normal numbers of ciliated cells (Fig. 8 C). Interestingly, midline cilia can no longer be detected in *Tmem107^schlei^* embryos later in development (6-7 somites stage, not shown). Notably, cilia are still absent from the definitive endoderm in *Tmem107^schlei^* embryos (Fig. 8 C), suggesting a differential requirement for TMEM107 function in primary cilia of distinct embryonic tissues, and that despite the enrichment for *Tmem107* transcript at the node, the definitive endoderm is more sensitive to the *Tmem107^schlei^* mutation.

While the presence of midline cilia correlated with normal situs and midline expression of *Lefty1* in *Tmem107^schlei^* embryos, these findings raised questions about what was occurring in *Mks1_krc_* embryos, which express midline *Lefty1*, but develop left isomerism. In the node, immunofluorescent analyses confirmed our previous SEM studies, revealing very few cilia in *Mks1_krc_* embryos (Fig. 8 D, E). Consistent with this, midline cilia were absent in *Mks1_krc_* embryos (Fig. 8 D, E). Together our findings suggest that loss of midline cilia drives left isomerism through the loss of midline barrier function in *Mks1^krc^*, *Tmem107^null^*, and potentially other cilia mutants, regardless of *Lefty1* expression.

**Figure 8.**
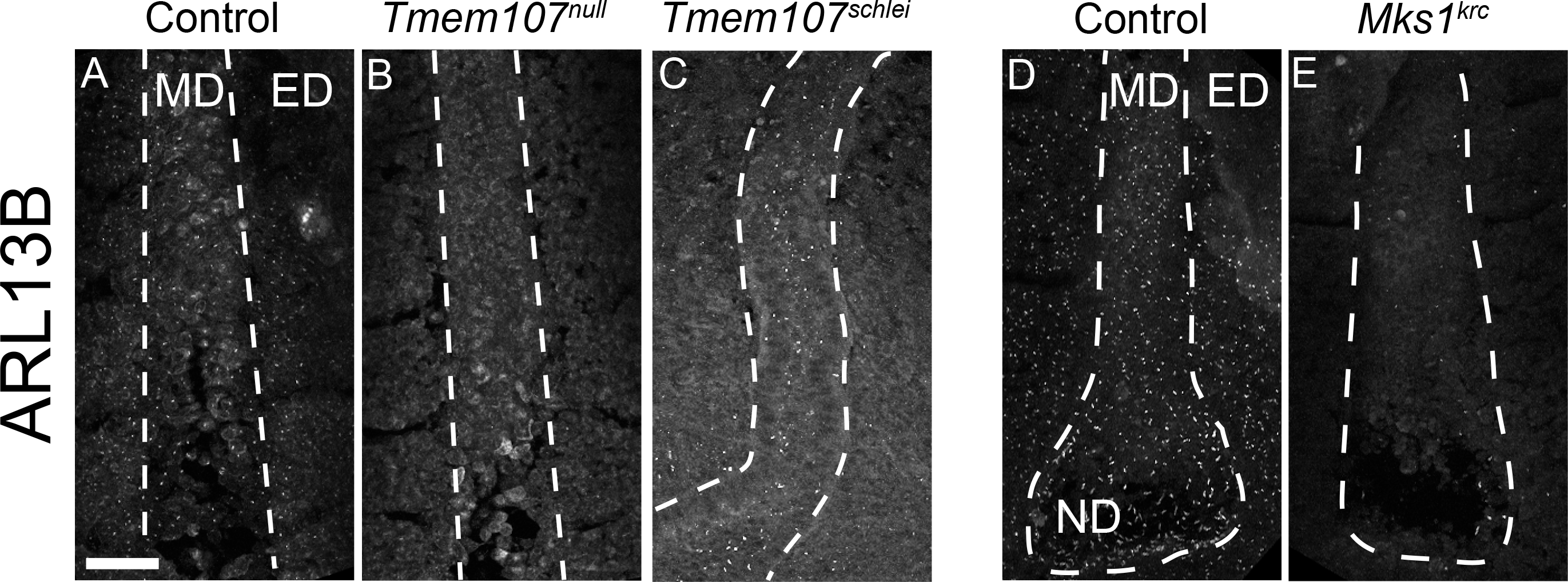
Absence of cilia in the midline correlates with the laterality defects. Maximum intensity projected Z-stacks of confocal immunofluorescent images of E8.0 midlines (A-C) and nodes with midlines (D, E) using ARL13B as a cilia marker. (A, D) Control embryos have numerous cilia in the midline (A, D), node (D) and the surrounding endoderm (A, D). (B) Only occasional cilia can be observed in the midlines of *Tmem107^null^* embryos. (C) A significant population of ARL13 B positive cilia is consistently observed in the midline of *Tmem107^schlei^* embryos. (E) Similar to *Tmem107^null^* embryos, *Mks1_krc_* embryos have very few cilia in the nodes (Weatherbee, *et al*, 2009), and only occasional cilia in the midline. Dashed lines outline the midline area. MD - midline, ED - endoderm, ND - node. Scale bar is 50μm.

## DISCUSSION

Node cilia have been a major focus of L-R patterning for their roles in generating the nodal flow, sensing that flow and leading to the original break in symmetry (McGrath, Somlo, Makova, *et al*, 2003; Nakamura, Saito, Kawasumi, *et al*, 2012; Schweickert, Vick, Getwan, *et al*, 2010; Shinohara, Kawasumi, Takamatsu, *et al*, 2012; Yoshiba, Shiratori, Kuo, *et al*, 2012). However, our understanding of how the resultant asymmetry is maintained, once initiated, is less clear. In this study we uncovered a distinct role for midline cilia in establishing the midline barrier during L-R patterning. The *Tmem107^null^* mutant has a discrepancy between abnormal, randomized L-R patterning markers at the node, and left isomerism at later stages. We traced the cause of this discrepancy to the loss of the midline barrier formation, and loss of midline *Lefty1* expression. Strikingly, we found that *Lefty1* can be expressed in cilia mutants that have lost midline SHH signaling, however its expression is not sufficient to prevent left isomerism. Instead, absence of midline cilia, regardless of *Lefty1* expression, correlates with bilateral *Nodal* expression and left isomerism. These findings indicate that cilia in the midline are required to maintain L-R asymmetry.

*Tmem107^null^* mutants have a nearly complete loss of cilia, with only a few visible in the node and the midline. Strong bilateral and often randomized *Cerl2* expression in the node results in weak and randomized *Nodal* expression, pointing to defects in generating or sensing flow. We see a further disconnect between the randomized signaling at the node and bilateral *Nodal* in the LPM. Although at early stages randomized *Nodal* at the node leads to its randomized expression in the LPM in a consistent manner, failure of the midline barrier allows Nodal to cross the midline and induce its expression in the opposite LPM, always leading to its bilateral activation at later stages and subsequent left isomerism. Together, the phenotypes observed in the *Tmem107^null^* mutant highlights the multiple roles for cilia during L-R patterning - initiating L-R asymmetry by establishing nodal flow, sensing the flow and ultimately maintaining L-R asymmetry through upholding the midline barrier functions.

Bilateral expression of *Nodal* is also the prevailing phenotype observed in mutants with a complete loss of cilia, suggesting that the cilia observed in *Tmem107^null^* mutants are effectively non-functional in the context of L-R patterning (Murcia, Richards, Yoder, *et al*, 2000; Takeda, Yonekawa, Tanaka, *et al*, 1999). However, some ciliary functionality is maintained in the embryo, since the mutants in our study survive until at least E16.5, while mutants with a complete loss of cilia, like *Kif3a* and *IFT88* die before E11.5 (Murcia, Richards, Yoder, *et al*, 2000; Takeda, Yonekawa, Tanaka, *et al*, 1999). Several studies have re-introduced cilia into the crown cells of cilia-deficient embryos, and have shown normal situs and proper midline barrier (Botilde, Yoshiba, Shinohara, *et al*, 2013; Yoshiba, Shiratori, Kuo, *et al*, 2012). Our interpretation of those results is that full recovery in those mutants is achieved due to the migration of ciliated cells from the node into the midline, as previously observed in wild-type embryos (Sulik, Dehart, Iangaki, *et al*, 1994). *Tmem107^schlei^* mutants contrast *Tmem107^null^* in that they don’t have L-R patterning defects, consistent with normal expression of all *Nodal* cascade markers examined, including *Lefty1*. Interestingly, *Tmem107^schlei^* has a similar strong reduction of cilia in the endoderm, as *Tmem107^null^*, but a near-normal numbers of cilia in the node, and numerous cilia in the midline. This indicates that *Tmem107*, and cilia in general, may be differentially required in different embryonic tissues, suggesting that despite being ubiquitous organelles, primary cilia likely have variable protein composition across tissue types.

MKS1 is a component of the transition zone complex that physically interacts with TMEM107, and the *Mks1_krc_* mutant shares many of the L-R characteristics observed in the *Tmem107^null^* mutant. Few cilia are present in *Mks1_krc_* E8.0 embryos, including the node and the midline. The embryos express *Pitx2* bilaterally, suggestive of bilateral activation of the Nodal cascade, but surprisingly, we find that many *Mks1_krc_* embryos have *Lefty1* in the midline. This is the first mutant, to our knowledge, with a near-compete loss of cilia, that expresses *Lefty1* in the midline. Yet, despite the presence of *Lefty1*, midline barrier function is lost. These data suggest that *Lefty1* expression is not sufficient to establish the midline barrier. We find that both *Tmem107^null^* and *Mks1_krc_* mutants lack cilia, while *Tmem107^schlei^* embryos have numerous cilia in the midline. We conclude that cilia in the node are required for the establishment of the asymmetry, and the midline cilia are required for the establishment of the midline barrier and maintenance of L-R asymmetry.

Cilia are hubs for *Shh* signaling in mammals and *Shh* is necessary for the proper formation of the notochord and the floor plate (Chiang, Litingtung, Lee, *et al*, 1996). Since *Lefty1* is expressed in the PFP, we expected its expression to be dependent on presence of cilia and *Shh* signaling (Tsiairis and McMahon, 2009; Tsukui, Capdevila, Tamura, *et al*, 1999). All three mutants that we describe show a strong reduction of *FoxA2* in their midlines, suggesting defective *Shh* signaling, but *Lefty1* is present in the midlines of *Tmem107^schlei^* and *Mks^krc^*. Additionally, loss of cilia in the midline of *Mks^krc^* mutants, coupled with presence of *Lefty1* expression, suggests that *Lefty1* expression may be independent of *Shh* and presence of cilia. Lack of *Lefty1* in the *Shh* and *FoxA2* mutant mice is likely due to structural defects and lack of the floor plate, rather than the failure to induce its expression through the SHH signaling cascade (Ang and Rossant, 1994; Dufort, *et al*, 1998; Tsukui, Capdevila, Tamura, *et al*, 1999; Weinstein, *et al*, 1994). Alternatively, it is possible that the low residual SHH signaling in these mutants is sufficient for *Lefty1* expression (Christopher, Wang, Kong, *et al*, 2012; Weatherbee, Niswander and Anderson, 2009).

In summary, this study shows that while *Lefty1* expression has previously been shown to be necessary for the establishment of the midline barrier, it is not sufficient to confer barrier function, and that cilia are required in the midline to establish the midline barrier. Loss of normal *Shh* signaling in the *Tmem107^schlei^* midline still allows for the formation of the midline barrier. Therefore, we propose a novel *Shh*-independent function for cilia in coordinating *Lefty1* expression and establishing the midline barrier.

## MATERIALS AND METHODS

### Mouse strains

Mouse experiments were performed in accordance with Yale Institutional Animal Care and Use Committee guidelines. The *Tmem107^tm1Lex^* mouse line, herein referred to as *Tmem107^null^*, has been previously described (Christopher, Wang, Kong, *et al*, 2012; Tang, Li, Tang, *et al*, 2010). The mouse line was initially generated on the C57BL/6N background, and the analysis in this paper has been performed on a mixed C57BL/6J;C57BL/6N background. *Mks1_krc_* (Weatherbee, Niswander and Anderson, 2009) and *Tmem107^schlei^* (Christopher, Wang, Kong, *et al*, 2012) mutant lines were crossed for at least 7 generations onto the C57BL/6J background prior to characterization. *Dnah11^GFP^* (*Lrd^GFP^*) (McGrath, Somlo, Makova, *et al*, 2003) mice were a gift of Martina Brueckner, and are maintained on a mixed genetic background. Genotyping primers and strategies are available in Supplementary Material, Table S1.

Timed matings were set up by pairing up male and female mice in the evenings. The following morning females were checked for the presence of vaginal plugs. If a plug was detected, noon on that day was designated embryonic day E0.5. Females were euthanized, and embryos were collected at E8.0-E16.5. Embryos were staged morphologically as E8.0 starting at the late head fold, and as E8.5 once they had 6 somites, and the heart had begun to loop.

### Expression analysis

*In situ* hybridization analyses were performed using standard methods (Nagy, 2003). Antisense probes were generated from the following cDNA plasmids: *Nodal* (Elizabeth Robertson), *Cerl2* (Martina Brueckner), *Lefty1*(Hiroshi Hamada), *Pitx2* (Axel Schweickart), *Ptch1, Shh* and *FoxA2* (Andrew McMahon).

The *Tmem107* riboprobe plasmid was generated by RT-PCR from C57BL/6J RNA with subsequent cloning into pCRII-TOPO and contains full-length mouse *Tmem107* (NM_025838.2) coding sequence. *In situ* hybridization with *Tmem107* sense probe showed no visible staining (not shown). The *Mks1* riboprobe plasmid was generated by RT-PCR from E10.5 whole embryo C57BL/6J RNA, with subsequent cloning into pBluescriptSK+ and contains mouse full-length *Mks1* coding sequence. Due to the large size of the *Mks1* transcript, *Mks1* riboprobe was hydrolyzed prior to use.

### Wholemount Immunostaining

Embryos were collected at E8.0-E8.5 and fixed in 4% paraformaldehyde (PFA) for 20 minutes at room temperature. Embryos were washed 3 times (15 minutes each) in phosphate buffered saline (PBS), then blocked with 1% goat serum and 0.1% Triton-X in PBS (PBGT) for 1 hour at room temperature. Primary antibodies in PBGT were added and incubated overnight at 4°C. After 3 15 minute washes in PBGT, secondary antibodies in PBGT were added for 2 hours at room temperature. Then, samples were washed 3 times (15 minutes each) in PBS and 0.1% Triton-X, and mounted on slides (Denville) using ProLong Diamond Antifade Mountant (Life Technologies). Antibodies used are available in Supplementary Material, Table S2.

### Imaging and image analysis

Images of organs and post-*in situ* hybridization embryos were taken on a Zeiss SteREO Discovery microscope. Immunofluorescence images were captured using Leica SP5 inverted confocal microscope using Leica Application Software. Image analysis was performed in ImageJ (NIH).

## ACKNOWLEDGEMENTS

We would like to thank Kasey Christopher for the original observation of left-right patterning defects in *Tmem107^null^* animals; Martina Brueckner and Karel Liem for continued support of this project through sharing resources, as well as constructive discussions of the results and feedback on the manuscript; Emilie League, Davis Li and other members of the Weatherbee and Liem laboratories for feedback on experimental design, and manuscript preparation. This manuscript is partially based on a dissertation submitted to fulfill in part the requirements for the degree of Doctor of Philosophy, Yale University.

## COMPETING INTERSTS

The authors declare no competing or financial interests.

